# Pathogenic variations illuminate functional constraints in intrinsically disordered proteins

**DOI:** 10.1101/2025.05.01.651640

**Authors:** Norbert Deutsch, Gábor Erdős, Zsuzsanna Dosztányi

## Abstract

Intrinsically disordered regions (IDRs) play key roles in cellular signaling and regulation, yet their contribution to human disease remains poorly understood. Here we analyzed nearly one million ClinVar missense variants, focusing on those located within IDRs defined by curated and predicted annotations. We found that pathogenic variants were significantly enriched in short linear motifs (SLiMs) and disordered binding regions, consistent with their central functional importance. To extend these insights beyond existing annotations, we applied *AlphaMissense*, a deep-learning pathogenicity predictor, and uncovered localized “island-like” patterns of elevated pathogenicity within IDRs. Leveraging these signals, we developed a classifier to prioritize predicted ELM motifs (PEMs), revealing thousands of candidate functional sites linked to major disease classes, including neurological, cardiovascular, and cancer-associated genes. Case studies in POLK, FOXP2, and LMOD3 illustrate how this framework connects genetic variation to molecular mechanisms, providing a scalable route to interpret variants of uncertain significance and advancing our understanding of pathogenicity in the disordered proteome.

**Summary:** This study reveals how deep-learning pathogenicity predictions can uncover functional motifs within intrinsically disordered regions, providing a new framework for interpreting genetic variation in the disordered proteome.

## Introduction

Missense variants represent the most common form of genetic variation in humans, yet their clinical interpretation remains a major challenge. Despite advances in cataloging human variation, the majority of missense mutations remain classified as variants of uncertain significance (VUS), limiting their utility in diagnosis and precision medicine ^1–3^. Historically, efforts to understand the pathogenicity of missense variants have focused predominantly on globular protein domains, due to their stable structures and rich functional annotations. However, over one-third of the human proteome consists of intrinsically disordered regions (IDRs)—segments that lack stable three-dimensional structure yet play essential roles in signaling, regulation, and molecular interactions ^4–7^.

IDRs often mediate transient interactions through short linear motifs (SLiMs) that control binding, localization, and post-translational modification ^8–9^. Mutations in these motifs can disrupt or create new binding interfaces, rewiring interaction networks ^10–12^. IDRs also contribute to the assembly of biomolecular condensates through liquid–liquid phase separation (LLPS). The misregulation of this process can also drive diseases, including neurodegeneration and cancer ^13–14 15 16^. Notably, roughly 20% of cancer driver genes are enriched in IDR mutations, and these disordered segments are surprisingly conserved, underscoring their regulatory importance ^5–17^.

Interpreting variants within IDRs, however, remains a major hurdle for computational prediction. State-of-the-art variant effect predictors (VEPs) that perform well in structured domains show markedly reduced accuracy in disordered contexts ^18–19^. Most rely on features such as evolutionary conservation and stable three-dimensional structure, which are largely absent in IDRs. Consequently, VEPs display high specificity but low sensitivity, correctly identifying abundant benign variants while misclassifying many pathogenic ones. This limitation highlights that a single pathogenicity score cannot capture the functional complexity of disordered regions. One strategy is to incorporate disorder-specific features, such as LLPS propensity, to improve variant classification ^20^. Here, we take a complementary approach, asking whether general predictors such as AlphaMissense can be used to extract latent functional information and reveal patterns of constraint unique to IDRs.

Here, we systematically analyze missense variants within IDRs to uncover hidden signals of pathogenicity. We confirm that although IDRs contain a smaller fraction of known pathogenic mutations in ClinVar, these variants are significantly enriched in functional elements such as short linear motifs (SLiMs). We further show that AlphaMissense scores display a localized “island-like” peak of pathogenicity centered on SLiMs—a signature absent from structured domains. Leveraging this signal, we developed a classifier that identifies and prioritizes functional motifs (Predicted ELM Motifs, or PEMs) across the disordered proteome. This framework reveals novel disease-relevant sites in proteins such as POLK, FOXP2, and LMOD3, providing a scalable route to interpret genetic variation in the disordered proteome.

## Results

### Clinvar mutations in disordered regions

To assess the extent to which intrinsically disordered regions (IDRs) challenge current variant interpretation, we systematically analyzed nearly one million missense variants, including about 70,000 pathogenic mutations from the ClinVar database, collectively covering less than 8% of all amino acid positions in the human proteome (Fig. 1a). Our analysis, which used the “Combined Disorder” model (see Methods), found that disordered regions comprise 36.8% of the human proteome (Fig. 1b). While uncertain mutations were distributed similarly to the proteome as a whole, benign mutations showed a notable enrichment in disordered regions (Fig. 1c). In contrast, and consistent with previous reports ^21–22^, we confirmed that pathogenic mutations are underrepresented in IDRs, with only 8.0% of all cataloged pathogenic mutations located in these regions (Fig. 1c). In total, 5,467 pathogenic mutations (2,298 unique positions) were identified in IDRs, corresponding to 2,957 distinct position–disease pairs (Fig. 1c, Supplementary Data 1). The list of pathogenic mutations in IDRs is provided in Supplementary Data 1. This residue-level view is complemented by a gene-level analysis, which shows that 798 genes harbor pathogenic variants within their IDRs, including 244 genes in which such variants are found exclusively in disordered regions (Fig. 1d). This underrepresentation of pathogenic variants likely reflects both historical biases toward studying structured proteins and the unique functional constraints operating in IDRs.

**Fig. 1:**
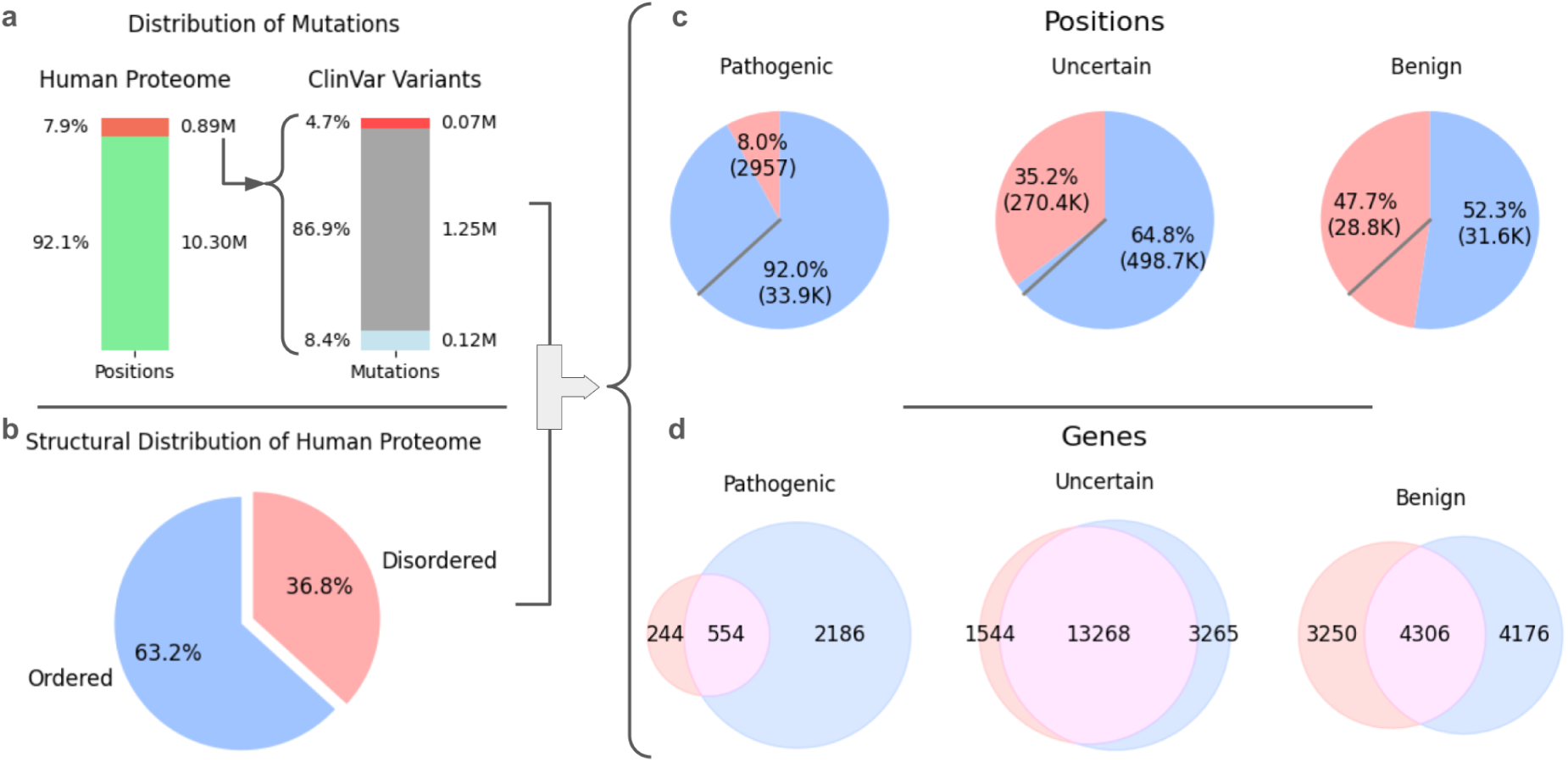
Distribution of ClinVar missense variants across ordered and disordered regions of the human proteome. **a** The left bar shows that of ∼10.3 million amino acid positions in the human proteome, 0.89 million (7.9%) are annotated with at least one ClinVar variant. The right bar shows the breakdown of all mapped ClinVar mutations by clinical significance : benign (blue), uncertain (grey), and pathogenic (red). **b** Structural composition of the human proteome, with 63.2% of residues classified as ordered (blue) and 36.8% as disordered (light red). **c** Positional distribution of ClinVar variants, stratified by structural state. Pie charts show the proportion of pathogenic, uncertain, and benign positions that fall within ordered (blue) vs. disordered (light red) regions. **d** Gene-level distribution of ClinVar variants. Venn diagrams show the number of genes harboring pathogenic, uncertain, or benign variants found exclusively in ordered regions (blue), exclusively in disordered regions (light red), or in both.

We examined the relationship between disease complexity and pathogenic mutations by classifying diseases as monogenic (single gene), multigenic (2–4 genes), or complex (>4 genes). Over 90% of ClinVar diseases involve fewer than five genes, yet nearly half of all mutations are linked to complex diseases and ∼20% to monogenic disorders. Mutations in intrinsically disordered regions (IDRs) showed a higher association with complex diseases than those in ordered regions, consistent with a role in signaling and regulatory pathways disrupted in these conditions (Fig. 2a). For each protein–disease pair, we assessed whether pathogenic variants preferentially affected disordered or ordered regions. When the majority (>60%) of mutations fell within intrinsically disordered regions, the disease was categorized as *disorder-specific*. This classification highlights disorders in which disordered segments are the main targets of pathogenic variation (Fig. 2b). Overall, 43.9% of mutations were disorder-specific, increasing to 50.5% for monogenic diseases (Fig. 2c).

**Fig. 2:**
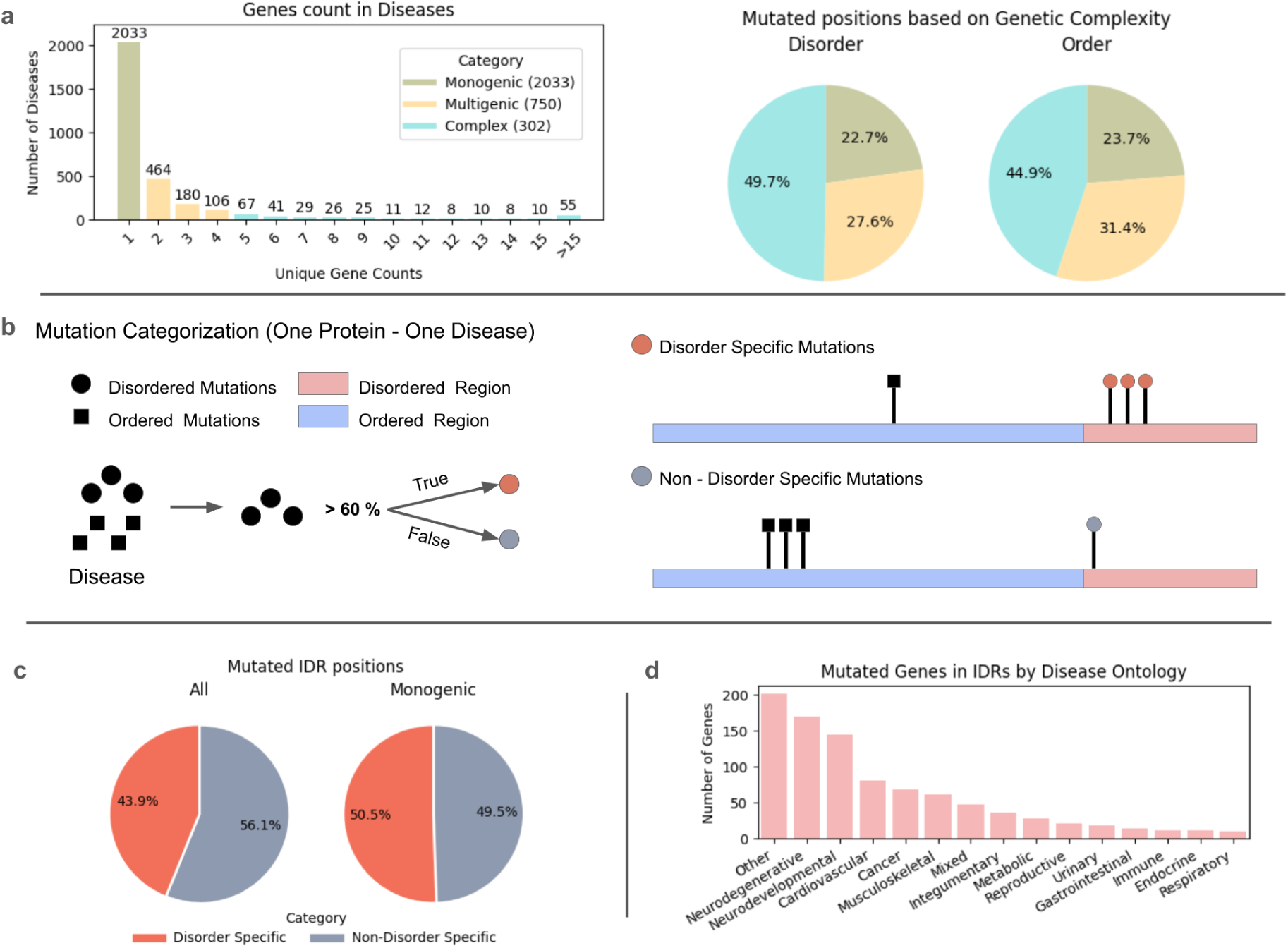
Genetic complexity and disease ontology of ClinVar variants. **a** Histogram illustrating the distribution of diseases by number of associated genes (left). Pie chart showing the distribution of pathogenic positions by genetic complexity. **b** Schematic illustration of the IDR mutation categorization. Proteins are classified as “Disorder Specific” when mutations predominantly occur within IDRs (> 60%), and “Non-Disorder Specific” otherwise. **c** Proportion of mutated IDR positions based on categorization. **d** Disease ontology for genes containing pathogenic IDR mutations (Others include diseases that could not be assigned to any specific disease category based on the disease ontology).

Diseases most impacted include neurodegenerative, neurodevelopmental, cardiovascular disorders, and cancer (Supplementary Fig. 1a). Mutations associated with these disease classes show a marked preference for intrinsically disordered regions (Supplementary Fig. 1b). Beyond these well-defined categories, IDR-specific mutations also occur in less characterized or multisystem disorders—such as developmental abnormalities—revealing their involvement in previously underappreciated disease types and underscoring the broad pathogenic relevance of disordered regions (Fig. 2d).

While these disease associations point to the functional importance of IDRs, the specific mechanisms often remain unknown. To investigate this, we first examined the available structural data from the PDB. This revealed that pathogenic variant sites have significantly greater experimental coverage than uncertain sites; however, structural information remains sparse for both, with 65% of pathogenic sites in disordered regions remaining uncharacterized (Figure 3a).

**Fig. 3:**
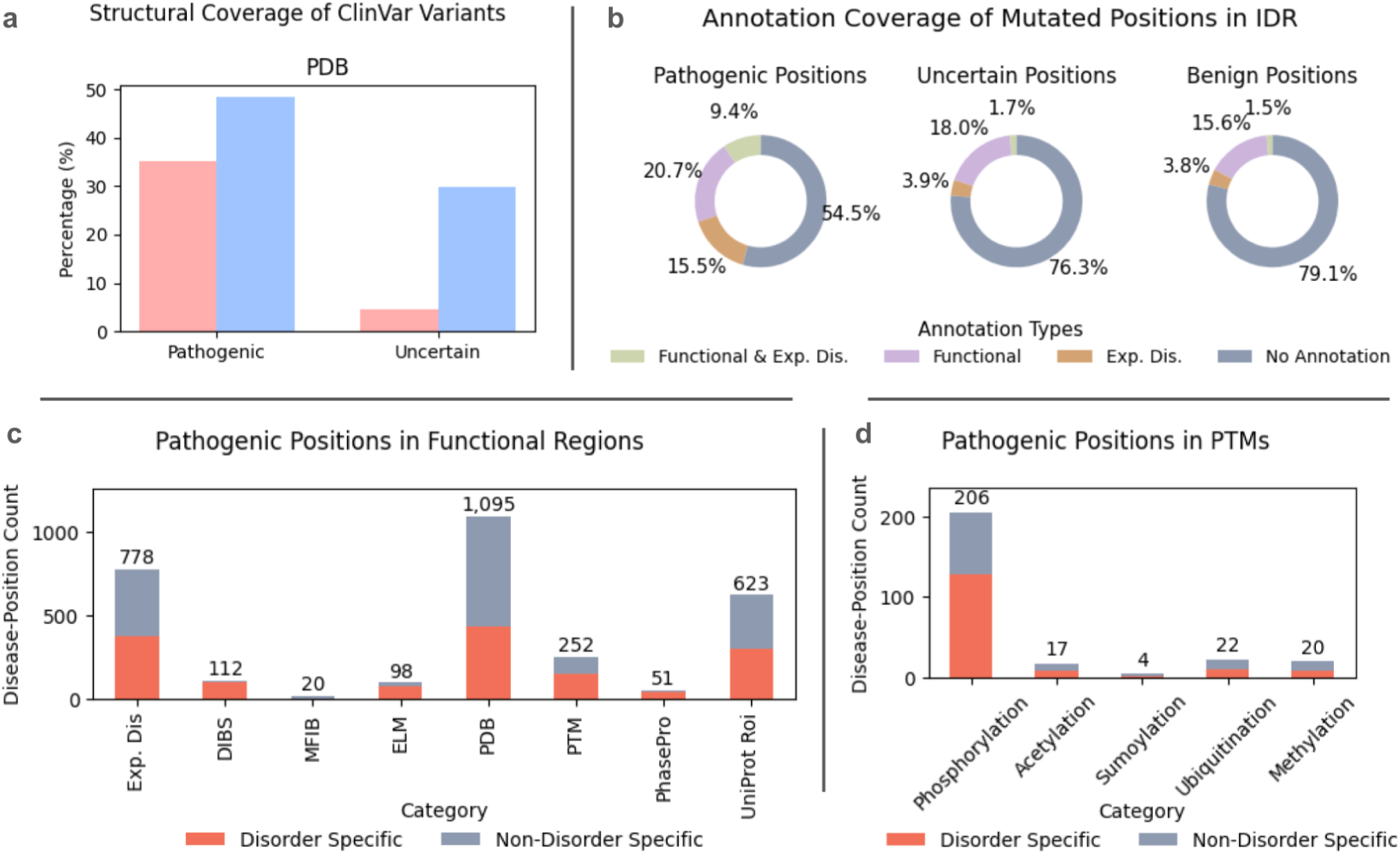
The Functional Annotation Landscape of ClinVar Variants. **a** Bar chart comparing the proportion of uncertain and pathogenic variant positions covered by experimental structures in the Protein Data Bank. **b** Donut plots showing the distribution of functional annotations among mutated residues within IDRs for pathogenic, uncertain, and benign variants. **c** Bar chart displaying the number of pathogenic positions overlapping specific functional annotation types, separated into disorder-specific and non-disorder-specific categories. **d** Bar chart showing the distribution of pathogenic variants by post-translational-modification (PTM) type, separated into disorder-specific and non-disorder-specific categories.

Next, we analyzed functional annotations compiled from UniProt and disorder-specific resources such as ELM^23^, DIBS^24^, MFIB^25^, and PhasePro^26^, together with post-translational modification (PTM) data from dbPTM^27^ and PhosphoSitePlus^28^. Pathogenic IDR variants were significantly enriched in known functional elements compared to uncertain and benign variants, including short linear motifs (SLiMs) and phosphorylation sites (Fig. 3c,d), yet 54.5% lacked any existing functional annotation (Fig. 3b). The functional constraint on these interaction sites was further supported by the distribution of benign variants: SLiMs from the ELM database showed the lowest benign mutation rate (0.94%) of all functional categories, consistent with strong purifying selection (Supplementary Fig. 1d). Together, these results indicate that although pathogenic variants preferentially occur in annotated functional elements such as SLiMs, most functionally important sites within IDRs remain unannotated. This gap underscores the limitations of current knowledge-based resources and motivates the development of predictive approaches to uncover uncharacterized functional regions.

### AlphaMissense as a predictive framework for identifying pathogenic and functional regions in IDRs

#### AlphaMissense captures functionally relevant mutations in disordered regions

To address the problem of unannotated intrinsically disordered regions (IDRs), we first evaluated the performance of AlphaMissense (AM). AM tended to classify all possible mutations in disordered regions as either fully pathogenic or fully benign, in contrast to the more gradual score distribution observed in ordered regions (Supplementary Fig. 2). Based on this observation, we assessed pathogenicity at the positional level by averaging AM scores across all 19 possible missense variants for each residue. Using this positional scoring approach, we found that while AM performed better on Clinvar variants at the variant level in structured regions, its balanced accuracy at the positional level was comparable between structured and disordered regions, with values of 76–77% (Fig. 4a).

**Fig 4.**
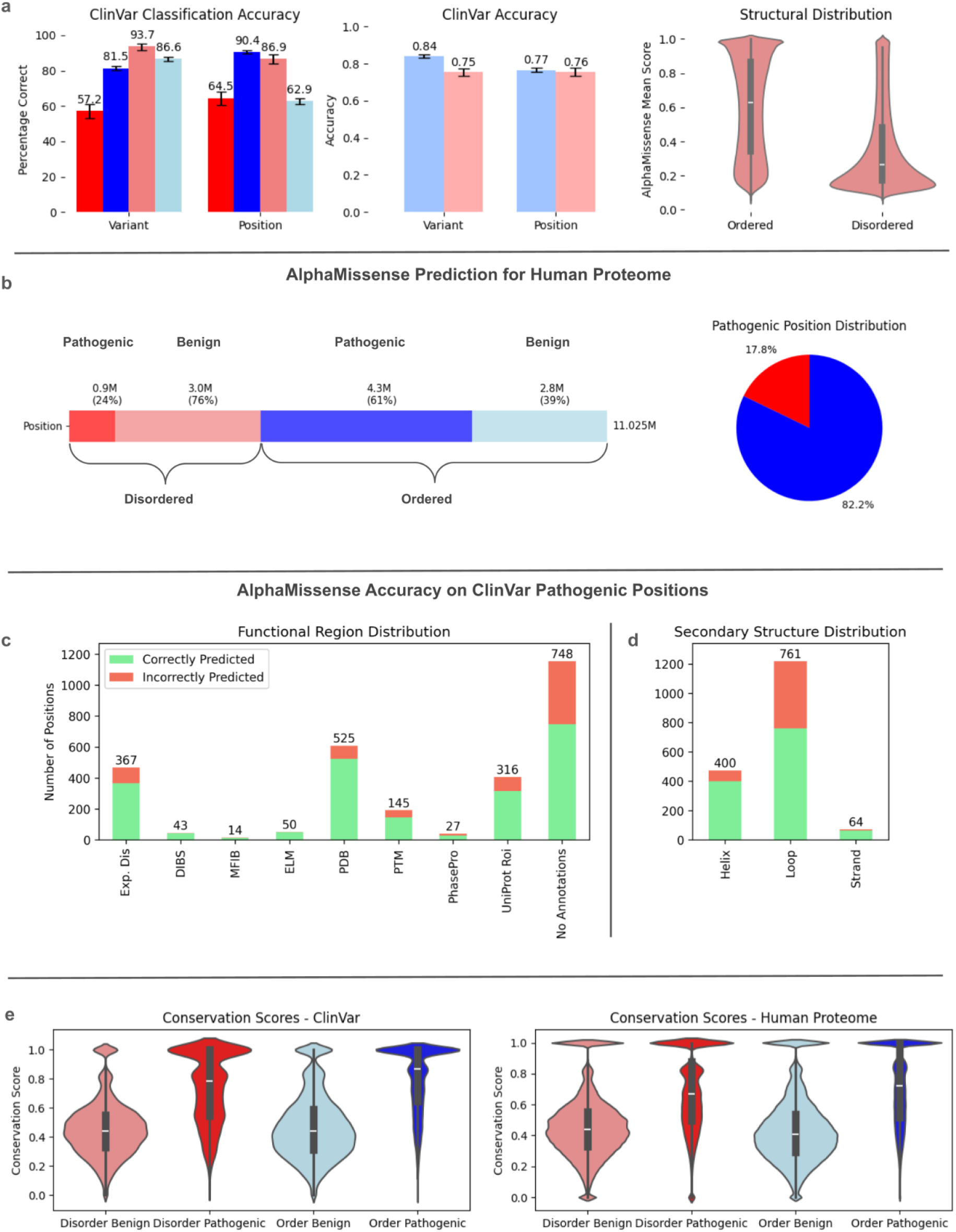
: AlphaMissense Performance, Proteome-Wide Prediction, and Correlation with Functional Features. **a** Bar charts showing AlphaMissense (AM) classification accuracy for ClinVar pathogenic (dark shades) and benign (light shades) variants in ordered (blue) and disordered (red) regions. The middle panel presents balanced-accuracy values from 1000 bootstrap iterations (75 % random sampling) at the variant and position levels, and the right panel displays violin plots of mean positional AM scores for ordered and disordered residues. **b** Bar chart illustrating the composition of predicted pathogenic (AM > 0.5) and benign residues in ordered and disordered regions of the human proteome, with a pie chart summarizing the proportion of predicted pathogenic residues within each structural context. **c** Bar charts reporting AM prediction outcomes for ClinVar pathogenic variants grouped by functional annotation, with green bars indicating correctly and red bars incorrectly predicted positions. **d** Bar charts showing AM prediction outcomes for ClinVar pathogenic IDR variants categorized by predicted secondary-structure elements, using the same color code as in (c). **e** Violin plots of evolutionary-conservation scores for benign and pathogenic variants from ClinVar (left) and for proteome-wide AM-classified residues (right).

Overall, mean pathogenicity scores were lower in disordered regions than in globular domains (0.36 vs. 0.60, respectively; Fig. 4a, right panel). Across the human proteome, AM predicted 4.7 million positions as pathogenic, but their distribution differed markedly between ordered and disordered regions (Fig. 4b). While 55% of positions in globular regions were predicted to be pathogenic, only 19% of IDR positions met this threshold. Despite this, IDRs contained a disproportionately high fraction (17.8%) of all predicted pathogenic positions—more than double the 8% currently annotated in ClinVar. These results suggest that a substantially larger fraction of IDR residues may have functional consequences than is currently recognized.

To better assess (AM) performance within IDRs, we stratified its accuracy according to existing functional and clinical annotation. AM showed the highest accuracy for variants located in regions under stronger functional constraint. Accuracy was significantly higher for mutations within annotated regions than for unannotated ones (75% vs. 62%), particularly for experimentally validated binding sites from DIBS, MFIB, and ELM (Fig. 4c). Similarly, disorder-specific mutations were predicted more accurately than non-disorder-specific ones (71% vs. 62%). The same trend was observed for disease complexity, with mutations associated with monogenic diseases showing the highest accuracy (73%), followed by complex (68%) and multigenic diseases (62%) (Supplementary Fig. 3).

To further examine the biological basis of AlphaMissense (AM) predictions in intrinsically disordered regions (IDRs), we analyzed their correlation with evolutionary conservation and predicted secondary structure. A strong correlation was observed with conservation: pathogenic variants in IDRs—both those reported in ClinVar and those predicted proteome-wide by AM—occur at significantly more conserved positions than benign variants (Fig. 4e). A similar pattern emerged for structural propensity, with IDR positions predicted as pathogenic by AM showing a higher tendency to form helices and strands in AlphaFold2 models (Supplementary Fig. 4).

This preference for structured elements also revealed a potential limitation. When AM was evaluated against known pathogenic variants located within IDRs, the misclassification rate was highest for mutations located in highly coil regions, whereas accuracy was greater for those in helices and strands (Fig. 4d). Altogether, these results support the functional relevance of pathogenic sites predicted by AM within IDRs, particularly those that are evolutionarily conserved and likely to adopt secondary structure elements.

#### AlphaMissense captures functional specificity and enables motif discovery in IDRs

While functionally annotated regions generally showed higher AlphaMissense (AM) scores, short linear motifs (SLiMs) exhibited a uniquely powerful signal. Binding regions within intrinsically disordered regions (IDRs) showed sharply elevated AM scores relative to their immediate flanking sequences, forming localized “island-like” patterns of predicted pathogenicity that were not observed in structured domains (Fig. 5a). This local contrast was strongest for annotated ELM motifs, with a mean score difference of +0.17 between the motif core and its surrounding residues (Fig. 5b). The signal also captured fine-grained variation within motifs: key functional residues exhibited significantly higher scores than non-key residues, consistent with their role in mediating specific interactions (Supplementary Fig. 5a). This pronounced, localized pattern was most evident for high-impact motif classes involved in degradation, docking, and targeting (Supplementary Fig. 6a), supporting its value as a robust indicator of functionally constrained sites.

**Fig. 5:**
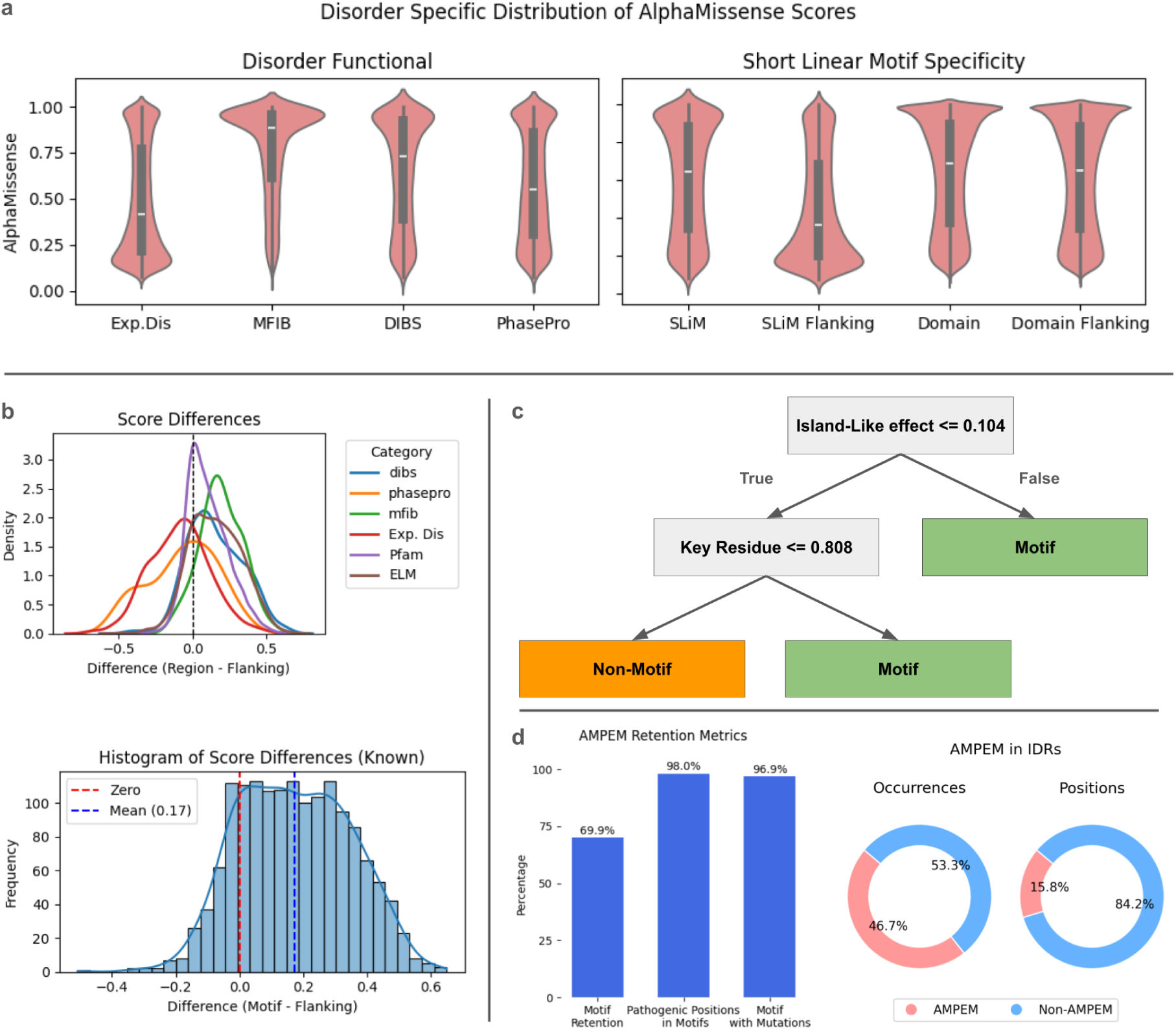
Predicting short linear motifs (SLiMs) using AlphaMissense scores. **a** Violin plots showing AlphaMissense score distributions for different functional categories within intrinsically disordered regions (IDRs) (left) and for SLiMs, PFAM domains, and their flanking regions (right). Sequential context is represented by windows of up to ten residues. **b** Density plots illustrating AlphaMissense score differences between annotated functional elements and adjacent flanking regions (top) and histogram of score differences for known motifs (bottom). **c** The decision tree model for the AMPEM (AlphaMissense-filtered Predicted ELM Motif) model, in which a region is classified as a true motif hit when the island-like score difference is ≥ 0.104 or the key-residue AlphaMissense score is ≥ 0.808. **d** Bar charts showing recovery rates for known ELM motifs, motifs containing pathogenic mutations, and mutated positions within motifs (left), and the fraction of regions and positions covered by predicted motifs (AMPEM) and by non-AMPEM regions (right).

Building on this finding, we evaluated whether AlphaMissense (AM) could extend beyond variant classification to support systematic identification of functional motifs across the disordered proteome. Due to their short and variable sequences, short linear motifs (SLiMs) are notoriously difficult to detect, and regex-based methods often yield numerous false positives. We used the sequence-pattern definitions from ELM, which yielded over one million predicted motifs (“Raw Predicted ELM Motifs,” RawPEMs). To filter these, we trained a simple decision tree classifier that leverages the AM signal. We chose a simple, interpretable decision tree as it performed comparably to more complex models while providing clear, rule-based logic for motif classification. Notably, the model selected just two features as most informative: (1) the “island-like effect” (the score difference between the motif and its flanks) and (2) the mean AM score of the motif’s key residues (**Figure 5c**).

The resulting set of motifs, termed AMPEMs (AlphaMissense-filtered Predicted ELM Motifs), represented a substantial refinement, reducing the initial pool by over 80% to ∼192,000 high-confidence predictions. This filtering effectively retained biologically meaningful sites. The classifier recovered 1,067 of 1,526 known motifs from the ELM database (69.9% recall) with a mean accuracy of 0.747 ± 0.040 (Supplementary Fig. 6b). It also preserved clinical relevance, retaining 97% of pathogenic positions (50 of 51) and 98% of disease-associated motifs (31 of 32) from the initial set (Fig. 5d; Supplementary Fig. 6c).

The proteome level analysis of the distribution of AMPEMs revealed that while nearly half of all IDRs (46.7%) contain at least one predicted motif, these motifs are sparsely distributed, covering only 15.8% of all disordered residues (Fig. 5d, right). This indicates that AMPEMs are not randomly distributed but tend to cluster within specific subregions, likely corresponding to functional hotspots. To generate a final dataset for clinical interpretation, we applied an additional filter to remove promiscuous, broadly matching ELM classes, yielding 8,654 CorePEMs (High-Confidence Predicted ELM Motifs). The CorePEM set retained a diverse range of motif types, with Ligand (LIG), Docking (DOC), and Degradation (DEG) classes being the most prevalent (Supplementary Fig. 5b,c).

We next conducted a series of tests to validate the functional relevance of the predicted motif sets. Using the highest-confidence CorePEM set, we first assessed signals of evolutionary constraint. Key residues within CorePEMs were under significantly stronger purifying selection, as indicated by a lower frequency of benign clinical variants compared with the initial regex-matched motifs (0.64% vs. 1.58%, *p* < 10⁻⁴⁰), confirming their functional constraint (Supplementary Fig. 6d). We then benchmarked the broader AMPEM set against independent datasets to evaluate its overall performance. AMPEMs recovered 87% of high-confidence motifs from the conservation-based tool SLiMPrints ^12–29^. Finally, we validated AMPEMs using a recent proteome-scale experimental dataset from peptide-phage display, which identified motifs where disease mutations disrupt protein binding ^12^. Our classifier correctly identified 59% of these experimentally validated functional motifs. Collectively, these results show that the classifier substantially reduces false positives from sequence-based searches while retaining a large proportion of biologically meaningful motifs.

### Prioritizing predicted ELM motifs for clinical interpretation of disease-associated mutations

Having established the biological relevance of the CorePEM dataset, we next prioritized motifs with potential clinical impact. Focusing on pathogenic and uncertain (VUS) variants from ClinVar, we selected motifs that showed a strong local enrichment of mutations, either a ≥3-fold enrichment of VUS relative to the rest of the protein, or containing variants found exclusively within the motif’s boundaries. This clinically driven prioritization yielded 228 HotspotPEMs, a curated set of mutation-enriched motifs. Among these, 67 HotspotPEMs were linked to pathogenic mutations (43 of which correspond to novel, unannotated motifs) and 161 were associated with VUS (132 novel), with most variants being disorder-specific (Fig. 6b, top panels). HotspotPEMs enriched in pathogenic mutations were mainly linked to neurodevelopmental and neurodegenerative conditions, with additional cases in the “Other” category representing rare monogenic disorders with developmental, immunological, or neuromuscular features. These findings highlight the broad clinical relevance of motif disruption within intrinsically disordered regions.

**Fig. 6:**
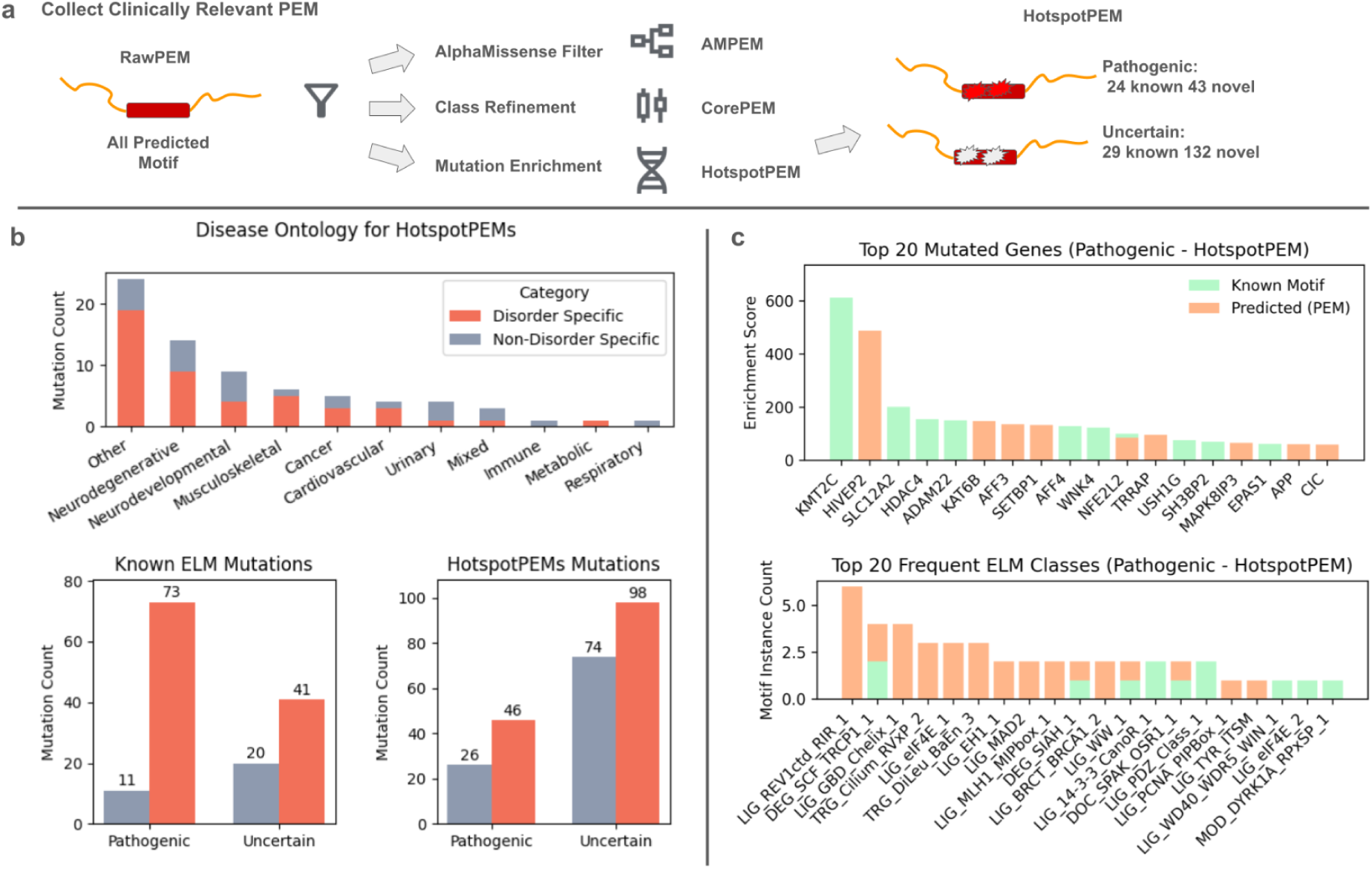
Identifying predicted motifs with potential disease relevance. **a** Workflow schematic illustrating the generation of HotspotPEM regions from clinically prioritized, mutation-enriched HotspotPEM, highlighting known and novel motifs with potential disease relevance. **b** Upper panel depicting the disease-ontology categories of pathogenic variants located within predicted motifs; lower panels showing the distribution of pathogenic and uncertain variants in known ELM motifs and in predicted HotspotPEMs. **c** Bar charts displaying genes with the highest numbers of pathogenic HotspotPEM mutations and the most frequent ELM motif classes, indicating known and newly predicted categories.

We next examined which specific genes and motif classes were recurrently targeted by pathogenic HotspotPEMs (Fig. 6c). Several genes—including *HIVEP2*, *KAT6B*, and *AFF3*—showed strong enrichment of pathogenic mutations within HotspotPEMs but did not overlap with previously annotated motifs, highlighting their potential clinical relevance. At the motif level, LIG-type motifs were the most frequently observed, although other motif classes were also represented. These findings underscore the functional diversity of disease-associated HotspotPEMs and identify specific classes of short linear motifs as recurrent targets of pathogenic variation in the disordered proteome.

To illustrate how this framework can generate mechanistic insights, we next present three case studies that highlight both known and novel motifs. These examples show how HotspotPEMs connect sequence-level variants to molecular dysfunction, providing plausible, data-driven hypotheses for disease mechanisms. For community use, the full set of prioritized HotspotPEMs and the complete CorePEM catalog are available as curated resources (Supplementary Data 2 and Supplementary Data 3).

### Mechanistic hypotheses for disease mutations generated by the HotspotPEM framework

#### A putative actin-binding motif in *LMOD3* suggests a mechanism for nemaline myopathy

Nemaline myopathy is a muscle disorder caused by pathogenic variants in *LMOD3*, a protein essential for organizing sarcomeric actin filaments ^30^. Our framework identified a HotspotPEM in the C-terminal disordered region of *LMOD3* (residues 535–553) corresponding to a previously unannotated but predicted actin-binding WH2 motif (LIG_Actin_WH2_1). This finding aligns closely with *LMOD3*’s established role in actin regulation.

Two known pathogenic mutations (R543L and L550F) occur directly within this predicted motif (Fig. 7a). We propose that these mutations impair *LMOD3*’s ability to bind and stabilize actin filaments, thereby disrupting sarcomere dynamics and contributing to disease. This case illustrates how the HotspotPEM framework can reveal plausible molecular mechanisms by pinpointing novel functional sites that link genotype to cellular phenotype, providing clear targets for experimental validation.

**Fig. 7:**
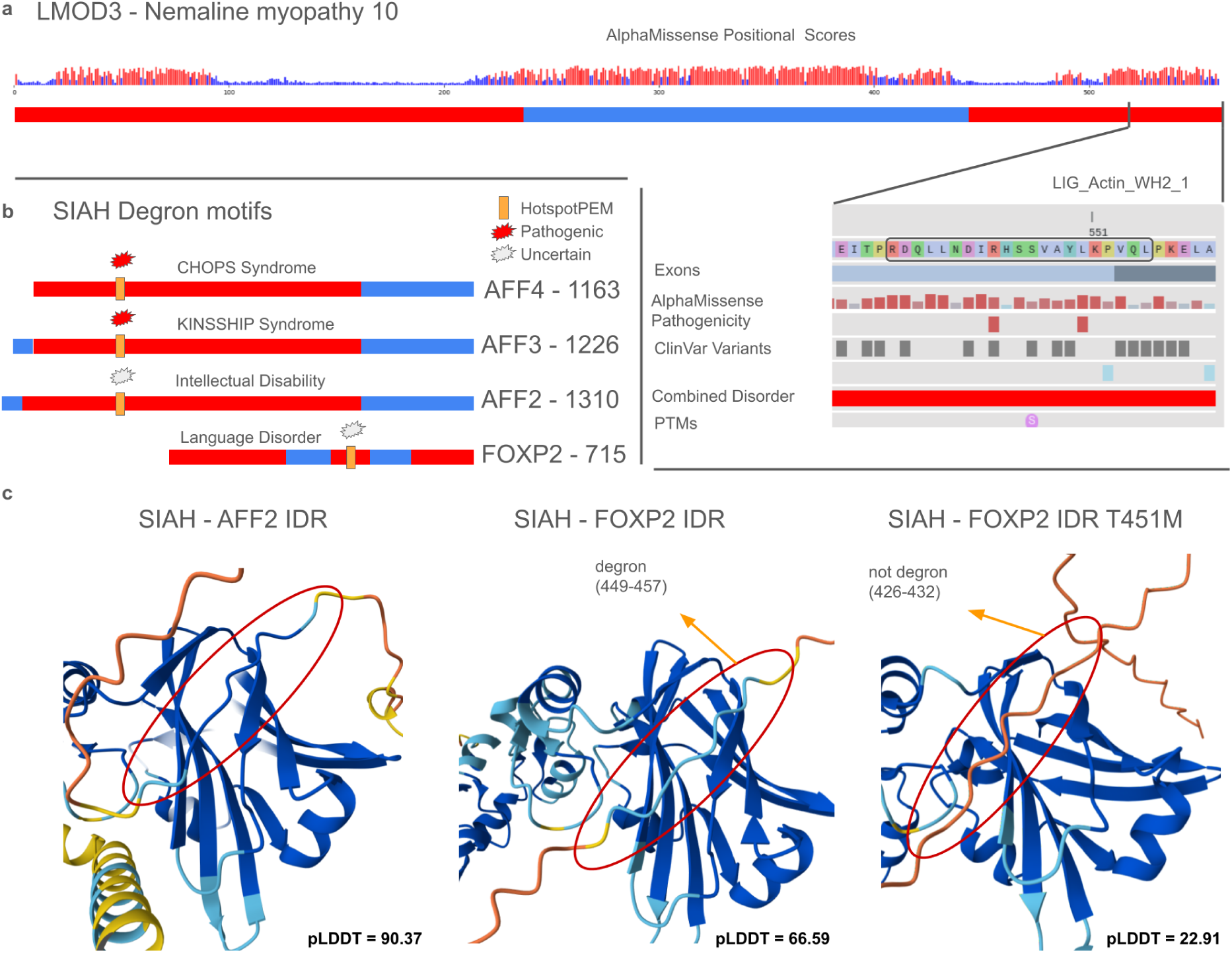
Functional motif disruption in intrinsically disordered regions (IDRs) associated with neuromuscular and neurodevelopmental diseases. **a** DisCanVis representation of the LMOD3 protein showing AlphaMissense positional scores and predicted ordered (blue) and disordered (red) regions. The highlighted segment shows the location of a predicted actin-binding WH2 motif overlapping two pathogenic variants (R543L and L550F) linked to nemaline myopathy. Tracks indicate exons, AlphaMissense pathogenicity, ClinVar variants, disorder annotations, and post-translational modifications. **b** Schematic overview of predicted SIAH degron motifs in AFF4, AFF3, AFF2, and FOXP2, highlighting associated pathogenic and uncertain ClinVar variants and their positions within disordered regions. The corresponding HotspotPEM regions and disease associations are indicated. **c** AlphaFold3 multimer models of SIAH bound to the IDR segments of AFF2 and FOXP2. Notably, the predicted degron region in FOXP2 (residues 449–457) interacts with SIAH in the wild-type model, whereas the T451M variant shifts the binding away from the degron and lowers model confidence (mean pLDDT scores shown below each motif).

### Predicted SIAH degron motifs point to dysregulated protein turnover in neurodevelopmental disorders

Our analysis identified multiple instances of predicted degron motifs recognized by the SIAH1 E3 ubiquitin ligase as HotspotPEMs, suggesting that impaired SIAH-mediated protein degradation may represent a shared mechanism across several disorders. Such motifs were enriched with pathogenic or uncertain variants in four key transcriptional regulators—*AFF3*, *AFF4*, *AFF2*, and *FOXP2*.

Pathogenic mutations in *AFF3* and *AFF4*, linked to KINSSHIP and CHOPS syndromes, respectively (Inoue *et al.*, 2023), cluster within the predicted degron sites. *AFF2*, another paralog, harbors uncertain variants at its predicted degron, suggesting that a related degradation mechanism may be perturbed in non-syndromic X-linked intellectual disability. Similarly, uncertain variants in *FOXP2*—a transcriptional repressor critical for cortical development and language—are concentrated within a predicted C-terminal SIAH degron (Fig. 7b). This leads to the testable hypothesis that these variants impair *FOXP2* degradation, resulting in its accumulation and prolonged repression of target genes.

To evaluate structural plausibility, we used AlphaFold to model the interactions. In all cases, the predicted degron motifs docked with high confidence into the SIAH1 binding interface (Fig. 7c). Introducing the *FOXP2* T451M variant in silico reduced model confidence at the degron site. Together, these findings provide a compelling, structurally supported hypothesis linking a shared molecular mechanism—disrupted protein turnover—to a group of related neurodevelopmental disorders.

### Predicted PCNA and REV1 interaction motifs provide a structural rationale for cancer mutations in disordered regions of POLK

The HotspotPEM framework offers a clear structural rationale for uncharacterized cancer mutations in the disordered regions of DNA polymerase kappa (POLK). While POLK’s role in translesion synthesis (TLS) and cancer is known, the functional impact of variants outside its catalytic domain has been ambiguous. Previous work had established a REV1-binding RIR motif (residues 565–576) but struggled to define a clear interaction with the key partner PCNA, proposing only a non-canonical C-terminal PIP motif that required additional residues for stable binding ^31^

We also identified a C-terminal RIR-like motif (residues 866–870) that overlaps the previously proposed PIP site. Modeling indicates that this motif preferentially binds REV1, forming an alternative interaction site distinct from the known RIR motif (Fig. 8c), and it continues to favor REV1 even in multimeric docking with both PCNA and REV1 (Fig. 8d). These results suggest that the newly identified PIP-like motif functions as the primary PCNA-binding site, whereas the C-terminal motif may serve as a secondary or context-dependent interface. The overlapping sequence characteristics and binding flexibility of PIP- and RIR-like motifs highlight the inherent ambiguity of current motif definitions in disordered regions, consistent with the view that these elements form part of a broader, promiscuous class of *PIP-like* motifs capable of engaging multiple partners involved in genome maintenance ^32^.

**Fig. 8:**
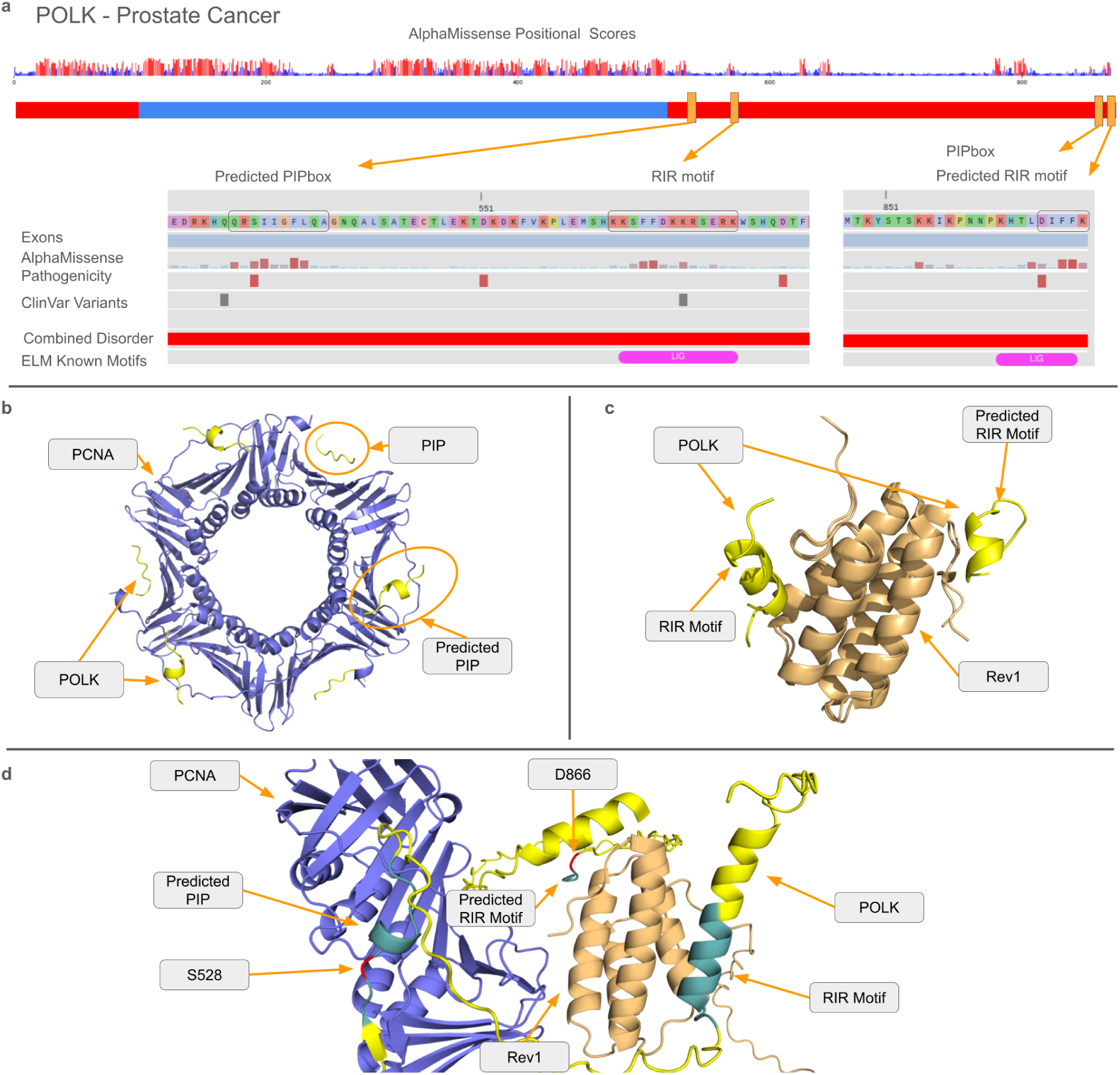
Cancer-linked motif disruption in disordered regions of POLK. **a** DisCanVis visualization of POLK showing AlphaMissense positional pathogenicity scores, disorder prediction, and ClinVar missense variants mapped to intrinsically disordered regions (IDRs). The figure highlights predicted PCNA-interacting (PIP) and Rev1-interacting (RIR) motifs with annotated sequence tracks. **b** AlphaFold3 multimer model of POLK bound to PCNA, displaying both annotated and predicted PIP-box motifs. **c** Structural alignment of the experimentally determined Rev1–POLK complex (PDB 4GK5) with the AlphaFold3 multimer model, illustrating the interaction of the Rev1 C-terminal domain with the known motif (residues 565–576) and the predicted RIR motif (residues 866–870). **d** Full multimeric AlphaFold3 model including POLK, PCNA, and the Rev1 C-terminal domain, showing predicted PIP and RIR motifs in yellow and pathogenic missense variants S528 and D866 in red.

Based on our findings, we propose a precise mechanistic hypothesis: the S528F mutation impairs PCNA docking at the primary PIP site, while the D866N mutation disrupts REV1-mediated recruitment. Both events likely compromise the assembly of the translesion synthesis (TLS) complex, promoting genomic instability. This case study demonstrates how our framework can move beyond ambiguous assignments to generate structurally-grounded hypotheses for previously uncharacterized cancer variants in disordered regions.

## Discussion

Intrinsically disordered regions (IDRs) comprise more than one-third of the human proteome yet remain markedly underrepresented in clinical variant annotations, accounting for only ∼8% of pathogenic mutations in ClinVar. This imbalance reflects a long-standing methodological bias: most pathogenicity studies have focused on folded domains, whose well-defined structures facilitate both experimental and computational interpretation. In contrast, the structural heterogeneity of IDRs makes it difficult to assess how mutations alter function, leaving a major gap in our understanding of disease mechanisms operating within the disordered proteome.

Our study addresses this gap by revealing that AlphaMissense (AM), a deep learning–based pathogenicity predictor, contains an untapped signal that can identify localized functional sites in disordered regions. Specifically, we discovered a characteristic “island-like” pattern in AM scores — sharp, localized peaks over short linear motifs (SLiMs) that sharply contrast with their surrounding sequence. These peaks mirror conservation-based signatures described in evolutionary studies ^29^ and indicate strong functional constraints. Strikingly, this signal emerges despite AM not being trained on ClinVar data or tailored to disordered proteins, suggesting that even general-purpose predictors implicitly encode information about functionally constrained sites in IDRs.

The significance of this finding is twofold. First, it provides a scalable route to detect functional elements in IDRs without relying on prior annotation or deep evolutionary conservation — two sources of information that are often sparse or misleading in these regions. Second, it complements other emerging frameworks for variant interpretation. While specialized predictors are being developed to evaluate the impact of variants on biophysical properties such as liquid–liquid phase separation (LLPS) ^20^, our approach captures a distinct mechanistic layer linked to compact, SLiM-mediated interactions. Similarly, it provides a computationally efficient alternative to structural modeling of domain–motif interfaces, offering a rapid, proteome-scale screen to prioritize candidate sites for experimental follow-up. We have previously shown that this “island-like” signal can also improve the prediction of binding regions in IDRs ^33^, underscoring its general utility.

Building on this principle, we developed a simple, interpretable decision-tree classifier that leverages these AM-derived profiles to filter over one million sequence-based motif predictions into a high-confidence set of *Predicted ELM Motifs* (PEMs). The resulting PEMs retain known pathogenic sites while dramatically reducing false positives, as supported by a significant depletion of benign variants in the final CorePEM set—consistent with purifying selection acting on these residues. Beyond the global statistics, the clinical relevance of PEMs is demonstrated through several representative case studies. In *POLK*, pathogenic mutations coincide with predicted PCNA- and REV1-binding motifs, providing mechanistic links to replication stress and cancer. Similarly, in *FOXP2* and *LMOD3*, disease-associated mutations map to predicted degron and actin-binding motifs, respectively, offering plausible explanations for the observed neurodevelopmental and muscular phenotypes. These examples illustrate how PEMs can connect individual sequence variants to functional disruption within the largely uncharted landscape of disordered regions.

While our computational framework is validated across independent datasets, we acknowledge its limitations. The most significant is the absence of direct experimental validation for novel predictions. The case studies presented here therefore represent testable hypotheses for future biochemical and cellular assays. Furthermore, our approach is inherently constrained by the current, imperfect definitions of SLiMs, which our classifier is designed to find. Consequently, the “island-like” signal may not capture all classes of SLiM-mediated interactions—particularly those that are weakly constrained, highly degenerate, or defined by features beyond a simple sequence pattern. Future work should integrate pathogenicity signals with complementary features such as conservation, physicochemical context, and phase-separation propensity to refine the map of functionally constrained elements in the disordered proteome.

In conclusion, our findings establish a new paradigm for variant interpretation in intrinsically disordered regions. By uncovering and exploiting a hidden functional signal within a general pathogenicity predictor, we provide an integrated and scalable framework for identifying and prioritizing pathogenic variants in the “dark proteome.” This approach not only expands our understanding of the molecular mechanisms underlying disordered protein function but also offers a practical avenue to reclassify variants of uncertain significance (VUS), a persistent bottleneck in genetic diagnostics.

## Methods

### Transcript mapping and annotation

We mapped GENCODE v44^34^ transcripts to the corresponding canonical protein sequences from UniProt^35^ (release 2024_01), following the procedure previously implemented in DisCanVis^36^. For all downstream analyses, we included only the main isoforms to ensure consistency across the dataset. All structural, functional, and pathogenicity annotations were assigned to these canonical sequences.

### Combined disorder model

To define the disordered state of protein positions, we combined experimental and computational data. Experimentally verified IDRs were obtained from MobiDB^37^ (categories: “homology-disorder-merge” and “curated-disorder-merge”). For proteins without experimental data, disorder was predicted using AlphaFold2-derived^38^ RSA values and IUPred3^39^ scores.

Pfam^40^ domains were mapped to the proteome, and each domain was labeled as disordered or ordered based on whether ≥50% of its residues were disordered. In disordered domains, all positions were considered disordered. In ordered domains, only terminal residues verified as disordered were retained. The final disorder state at each position was assigned hierarchically: experimental > AlphaFold2-RSA (>0.582) > IUPred3 (>0.4). Positions located in coiled-coil regions (DeepCoil^41^, threshold: 0.5) and collagen genes were excluded.

### Functional and structural annotations

Functional annotations were sourced from the second version of DisCanVis and mapped to Gencode v44. We focused on disordered-specific annotations, post-translational modifications (PTMs), and curated regions from UniProt. UniProt’s “Region of Interest” entries were filtered to exclude general structural descriptors.

Disordered binding regions were extracted from MFIB ^25^ and DIBS ^24^ databases; SLiMs were obtained from the ELM database ^23^. Phase-separating regions were derived from PhasePro^26^. PTM annotations were compiled from dbPTM ^27^ and PhosphoSitePlus ^28^. PDB entries were mapped to Gencode transcripts, and missing residues were excluded. Secondary structure assignments from AlphaFold2 were computed using the 4th version of DSSP ^42^.

### ClinVar variant processing

Missense variant data were obtained from ClinVar (VCFv4.1, release date: 2023-08-13, GRCh38) and mapped to Gencode v44 transcripts. Only non-synonymous single-nucleotide variants (SNVs) were retained, excluding those at the first codon position to avoid potential mapping ambiguities. Variants were filtered to include only those labeled as “Pathogenic,” “Likely pathogenic,” or “Benign,” with at least one ClinVar star rating.

Variants annotated as “Uncertain significance,” “Conflicting interpretations of pathogenicity,” or “Not provided” were grouped together as variants of uncertain significance (VUS) for downstream analyses.

### Disease ontology classification

Disease ontology assignments were extracted from the MONDO Disease Ontology database^43^. Diseases were classified into 16 major organ/system-specific categories, such as Cancer, Cardiovascular, Neurodevelopmental, and Musculoskeletal. If a disease matched multiple categories, it was assigned to an organ-specific group when possible; otherwise, it was classified as “Mixed.” Any disease with “Cancer” in its ontology path was prioritized into the Cancer group. Diseases without mapped terms were assigned to the “Unknown” category.

These groups were used to assess genetic complexity by classifying diseases as Monogenic (1 gene), Multigenic (2–4 genes), or Complex (>4 genes).

### AlphaMissense

AlphaMissense scores were downloaded from the original publication and mapped to the transcripts. Scores from the first codon position were excluded. A score threshold of 0.564 was used to classify positions as pathogenic based on the original paper.

To evaluate predictive accuracy, ClinVar variants were filtered to reduce redundancy using DIAMOND^44^ (40% sequence identity). Proteins with equal numbers of pathogenic and benign variants were selected. Variants were classified as pathogenic or benign using the 0.567 threshold. For positional scoring, AlphaMissense scores were averaged across residues, and a threshold of 0.5 was used for binary classification.

### Short linear motif prediction (PEM)

A custom motif prediction pipeline was developed using ELM regex patterns to scan the human proteome. For each motif instance, we extracted the following features:

- The mean AlphaMissense (AM) score for the motif
- The mean AM score for the sequential motif region (up to 10 amino acids)
- The mean AM scores for key residues versus non-key residues
- The maximum AM residue mean score within the motif region
- The difference between sequential and motif AM scores
- The difference between key and non-key AM scores

To capture potential “island-like” pathogenic zones, we computed the difference between the motif’s mean AM score and that of its sequential region, as well as the difference between key-residue and non-key-residue AM scores. We focused on ELM classes predominantly located within intrinsically disordered regions (IDRs) and retained only those motif instances for which all key metrics were available. Additionally, we required that at least 80% of the motif sequence fall within disordered regions, as defined by our Combined Disorder model. This filtering yielded 1.1 million predicted motifs and 1,523 known motif sets.

For classification, we treated known motifs as the positive set and predicted motifs as the negative set. We trained a decision tree using scikit-learn’s DecisionTreeClassifier (maximum depth of 3, entropy-based splitting, and balanced class weights). We then applied 10-fold cross-validation with 1,000 bootstrap samples, each time selecting a random subset of predicted motifs as the negative set. An 80/20 train-test split was employed during each iteration.

For motif prioritization, we calculated local variant density within each predicted motif and compared it to the background density across the entire protein. Motifs were retained if their density was at least threefold higher than the background or if variants were exclusive to the motif region.

### Motif validation datasets

#### Peptide interaction dataset validation

To evaluate the functional relevance of PEMs, we utilized a dataset from a recent study^12^, which included experimentally tested domain–peptide interactions. The dataset contains peptide pairs (wild-type and mutant) mapped to interaction domains, with annotations on whether the mutation affected binding. We selected peptides containing regex-defined SLiMs and mapped them to our reference proteome. Motifs were filtered based on disorder using our Combined Disorder model, and each instance was scored using our decision-tree–based PEM predictor to assess whether it would be classified as a motif.

#### SLiMPrint validation

We also validated our predictions using SLiMPrint, a conservation-based method for identifying candidate SLiMs. Human motif instances were downloaded and mapped to our proteome and filtered based on the Combined Disorder model. We evaluated all mapped SLiMs as well as a high-confidence subset annotated with “Good,” “Strong,” “Motif,” or “Ok” scores. Each motif was then passed through our PEM prediction pipeline to determine classification.

#### Benign mutation rate estimation

To assess selective constraint, we calculated the benign mutation rate in annotated and unannotated IDRs. This was restricted to proteins containing benign variants. The mutation rate was defined as the proportion of disordered positions harboring at least one benign variant in each functional category (e.g., ELM, MFIB, PhasePro, experimental IDRs, and unannotated IDRs). For PEMs, we applied the same approach focusing on refined PEM and all predicted key residues.

### Data Availability

The datasets used in this study are publicly available from the following sources:

- **GENCODE v44** transcript annotations were obtained from the GENCODE database: https://ftp.ebi.ac.uk/pub/databases/gencode/Gencode_human/release_44/
- **ClinVar missense variant data** (release 2023-08-13, GRCh38) was downloaded from the NCBI FTP archive: https://ftp.ncbi.nlm.nih.gov/pub/clinvar/vcf_GRCh38/archive_2.0/2023/clinvar_20230813.vcf.gz
- **AlphaMissense variant effect scores** were retrieved from the DeepMind repository: https://storage.googleapis.com/dm_alphamissense/AlphaMissense_aa_substitutions.tsv.gz
- **Short linear motif annotations (ELM)** were accessed via the ELM database: http://elm.eu.org/downloads.html#instances
- **MFIB** (Mutual Folding Induced by Binding) and **DIBS** (Disordered Binding Sites) datasets were downloaded from their respective databases: https://mfib.pbrg.hu/downloads.php https://dibs.enzim.ttk.mta.hu/downloads.php
- **Experimental Disorder regions** were collected from **MobiDB** (accessed on 2024-04-11): https://mobidb.org/statistics?proteome=UP000005640
- **Disease ontology information** was obtained from the **MONDO Disease Ontology** project: https://mondo.monarchinitiative.org/pages/download/
- Shown examples was extracted from the second version of **DisCanVis**: https://v2.discanvis.elte.hu

## Supporting information

Supplementary Data 1: Pathogenic Variants in IDRs

Supplementary Data 2: Variants in HotspotPEM

Supplementary Data 3: CorePEM dataset

## Supplementary Data

Supplementary Data 1: [Pathogenic Variants in IDRs](https://docs.google.com/spreadsheets/d/1jHiQ8LB0WyT3BuqaEzrHdscAU84bWU_GVXCkpYV8fhI/edit?usp=sharing)

Supplementary Data 2: [Variants in HotspotPEM](https://docs.google.com/spreadsheets/d/1zNYIAIdZRp0BMGZ_6s5JtmndJzupW4ar72kjePahLT0/edit?gid=1540509754#gid=1540509754)

Supplementary Data 3: [CorePEM dataset](https://docs.google.com/spreadsheets/d/1hk4Hy4uT-B-Eg-1_qCgFEQ3LM2OgyYwNDaCfe87aFUY/edit?gid=0#gid=0)

## Funding

N.D. acknowledges “Supported by the EKÖP-KDP-24 University Excellence Scholarship Program, the Cooperative Doctoral Program of the Ministry for Culture and Innovation, and the National Research, Development and Innovation Fund.”

E.G. acknowledges “Supported by the EKÖP-24 University Excellence Scholarship Program of the Ministry for Culture and Innovation, funded by the National Research, Development, and Innovation Fund.”

Z.D. acknowledges “HORIZON-MSCA-2023-SE - Grant Agreement 101182949 “IDPfun2”.

This work was funded by the European Union. “HORIZON WIDERA 2023 IDP2Biomed - Grant Agreement 101160233”.

## Supplementary Materials

**Supplementary Figure 1).**
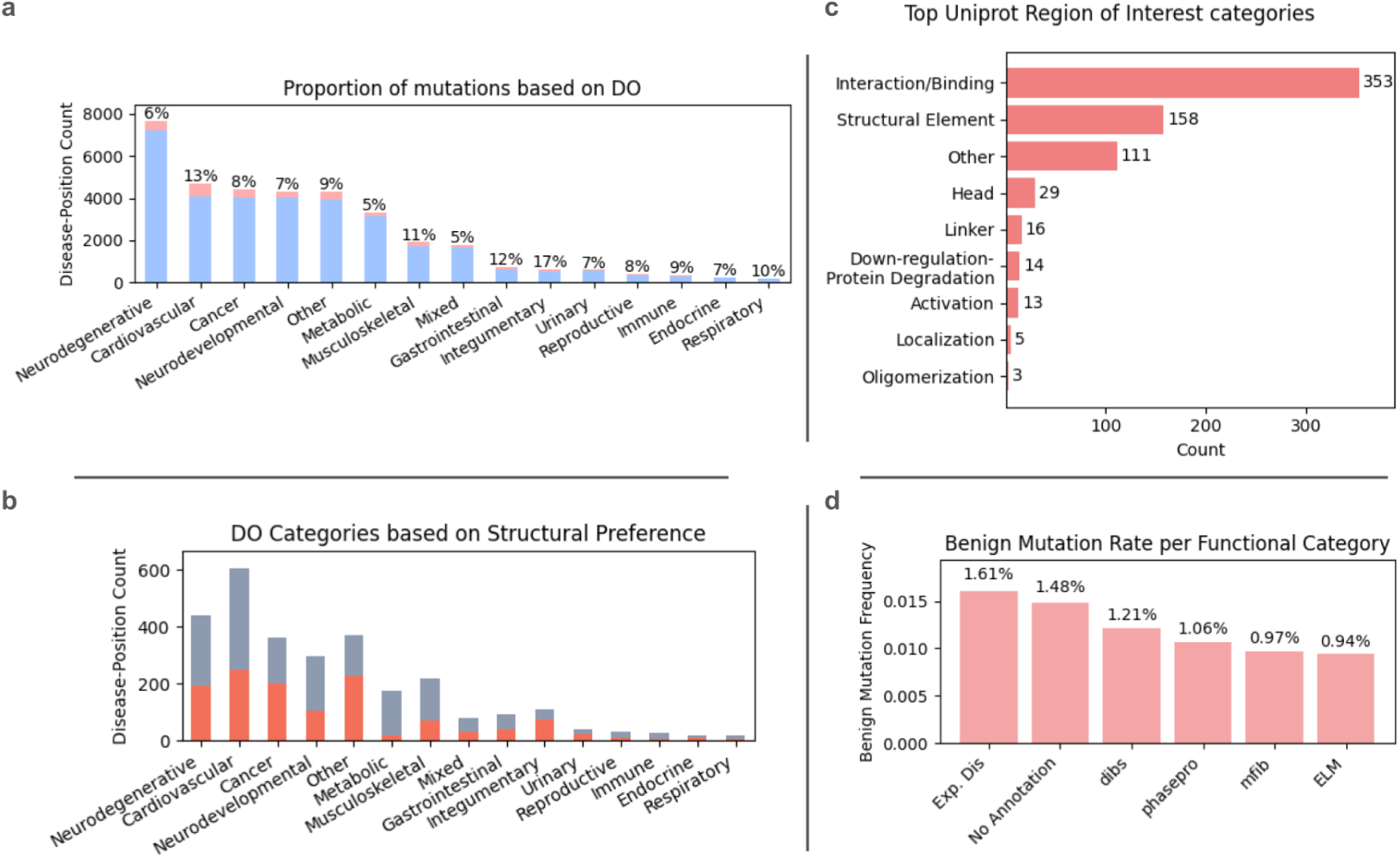
a) Structural classification of pathogenic positions by disease ontology. Blue indicates ordered regions, while light pink denotes disordered regions. The percentages represent the proportion of pathogenic positions located in disordered regions within each disease class. b) Distribution of pathogenic mutations within disordered regions across different disease ontology categories. c) Top UniProt region-of-interest (ROI) sites ranked by the number of pathogenic mutations. d) Benign mutation rates across different functional categories compared to unannotated disordered regions.

**Supplementary Figure 2).**
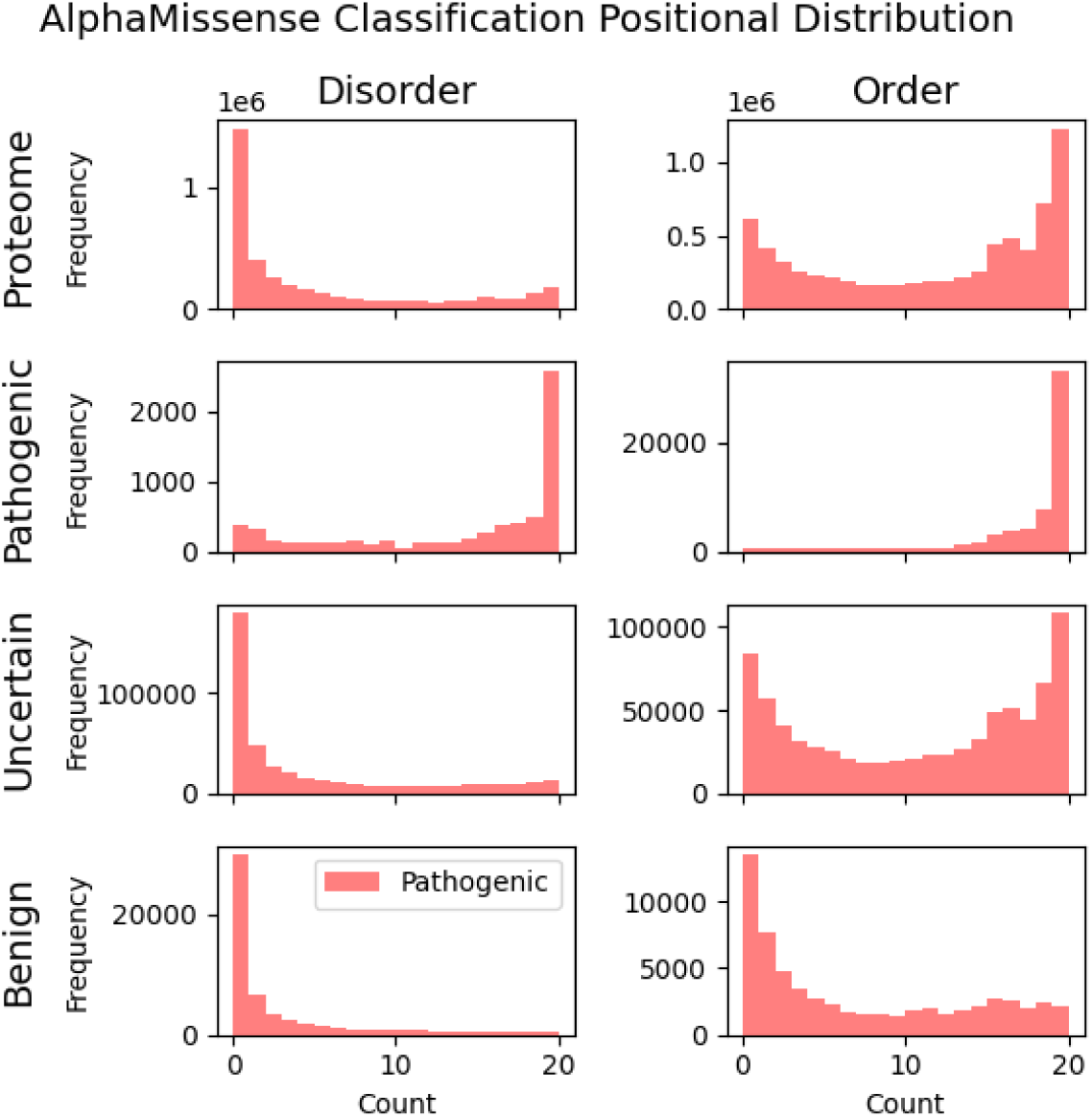
Distribution of AlphaMissense pathogenicity classifications for positions across the human proteome and for ClinVar-reported variants. Bars indicate the number of positions classified as pathogenic based on the mean AlphaMissense scores.

**Supplementary Figure 3).**
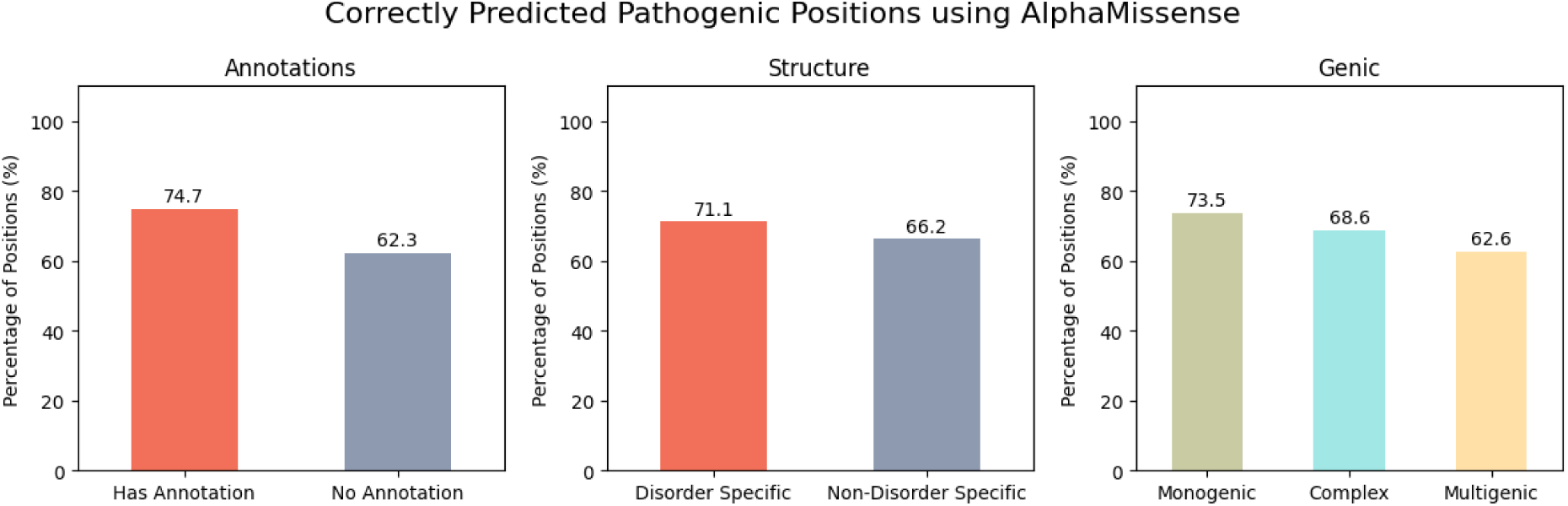
AlphaMissense Correctly Predicted Pathogenic Positions. In the top we present the accuracy based on annotations. In the bottom segment we show based on structural distribution and genic category.

**Supplementary Figure 4).**
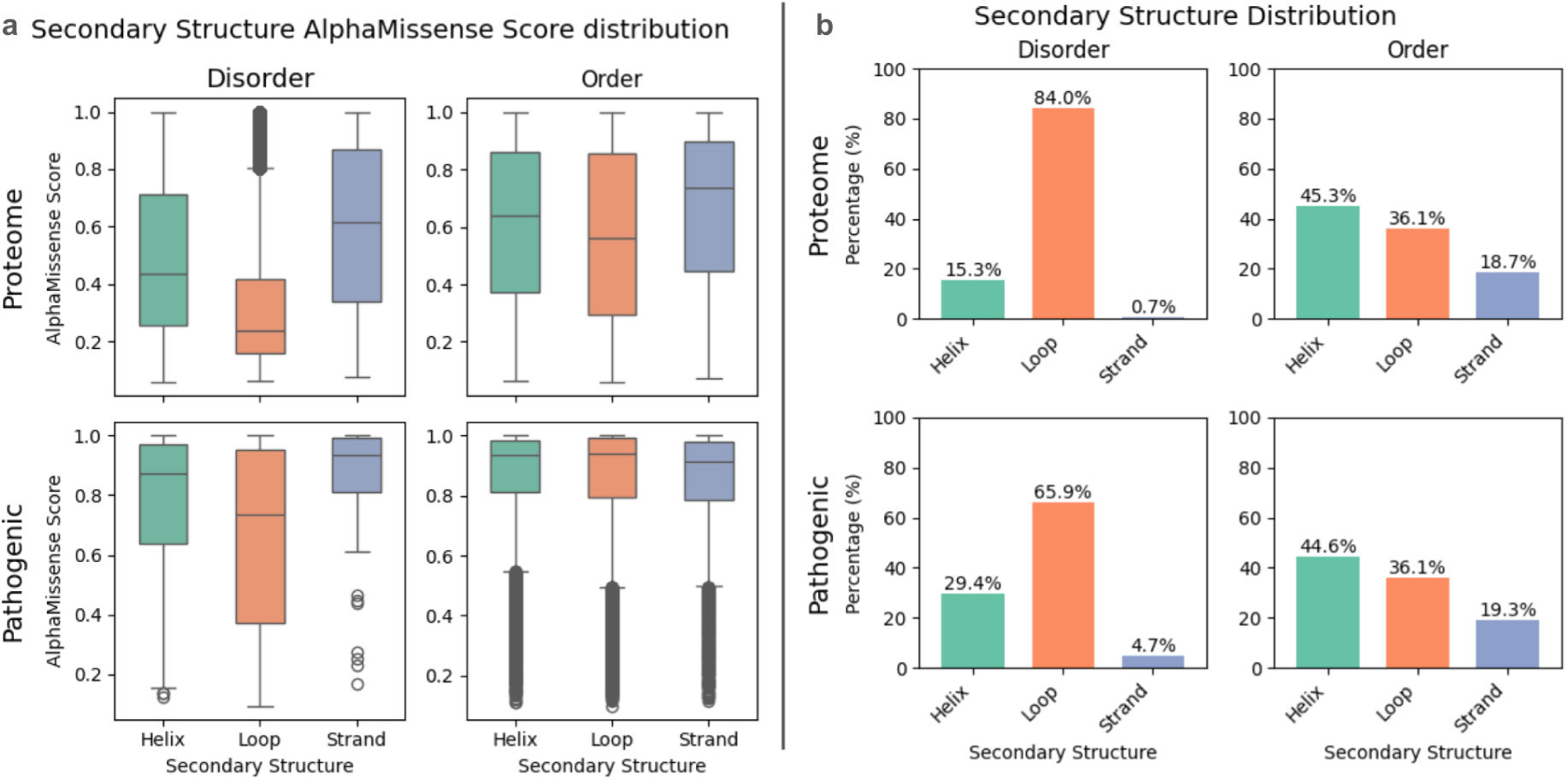
a) Distribution of Global AlphaMissense Scores for ClinVar and Human Proteome Positions b) Distribution for Secondary Structure for Proteome and ClinVar Pathogenic Positions.

**Supplementary Figure 5).**
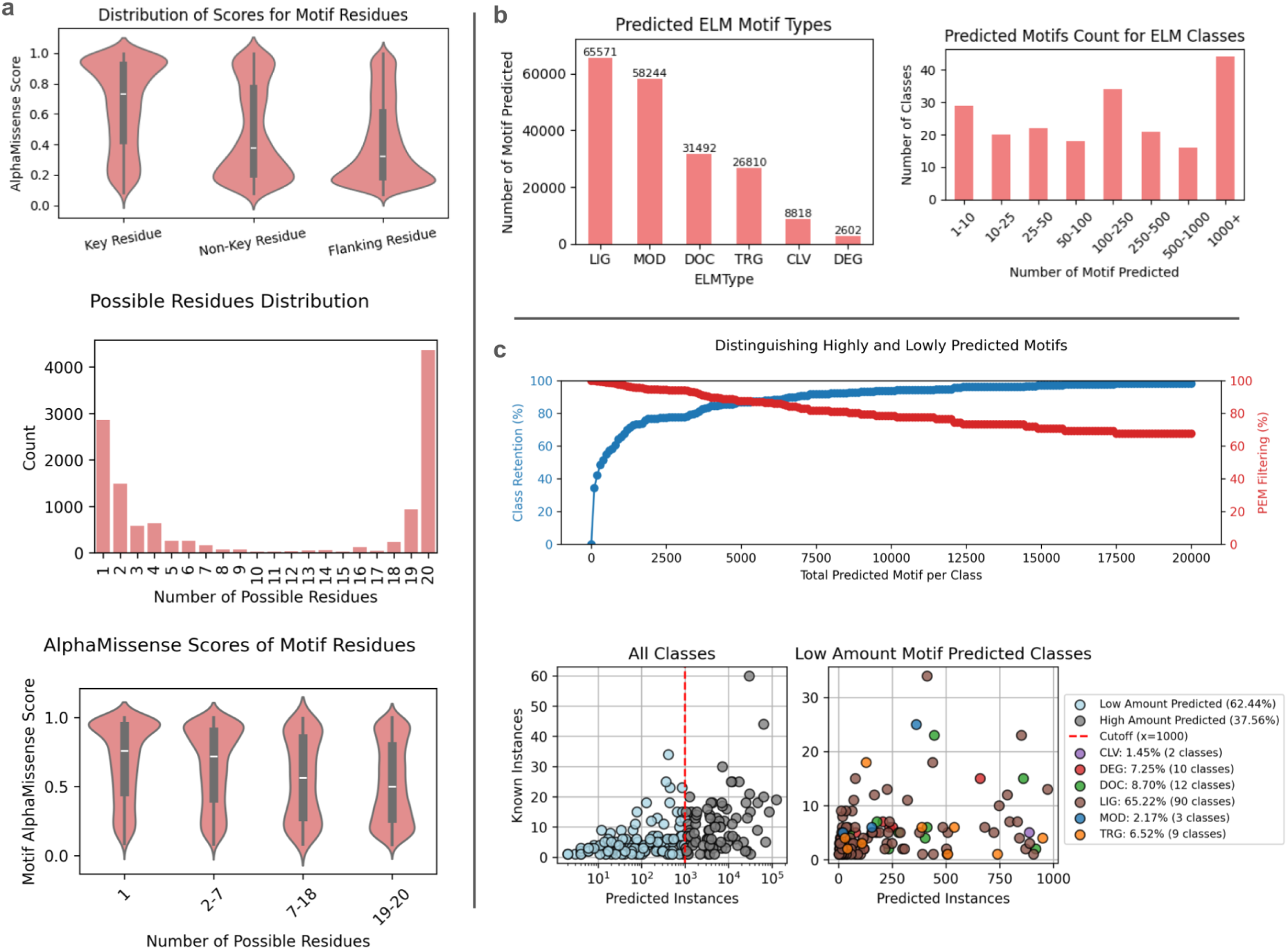
a) Distribution of AlphaMissense Scores for Motif Residues. Key residues (spanning 1–5 amino acids) within each motif display higher pathogenicity scores than both non-key residues (5–20 amino acids) and the surrounding flanking regions. Below we show a detailed distribution of the number of possible residues within known ELM motifs and the AlphaMissense score distribution of these residues. b) PEM prediction on human proteome. In the left figure we show the number of regions predicted for each protein. The right figure shows the motif length distribution for the predicted motif regions. c) Removal of ELM Classes with high Instance Prediction and the High Confidence Classes ELM Type Distribution. The bottom part contains predicted motif statistics. The left figure shows the ELM Type distribution of the predicted instances. The right figure shows the ELM classes’ predicted instances counts.

**Supplementary Figure 6).**
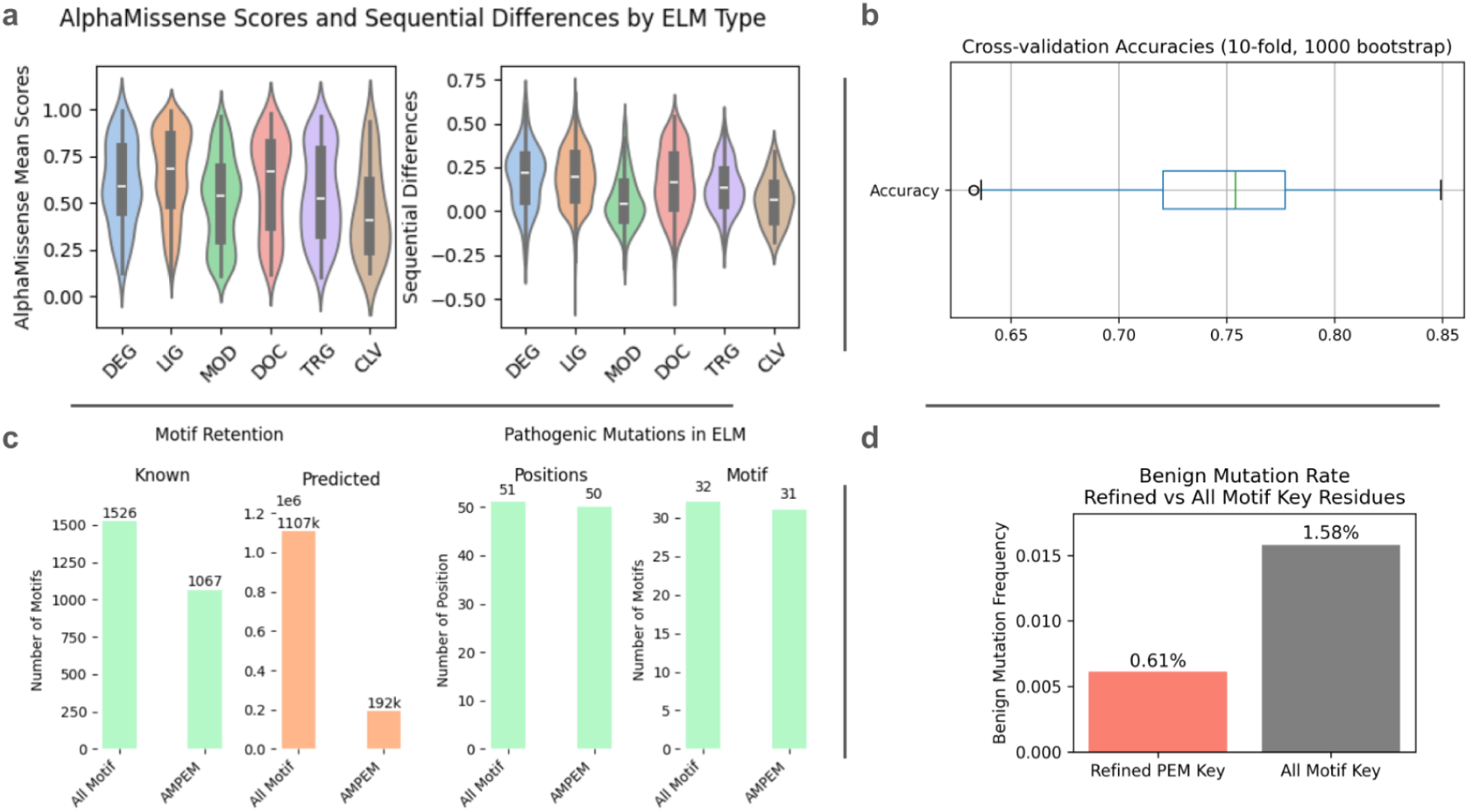
(a) AlphaMissense Mean Score Distribution for Various ELM Types and Differences Between Sequential and Motif Scores. Distinct SLiM types exhibit noticeable variation in AlphaMissense scores, as well as in their differences from the local sequence environment. Among these, LIG, DOC, and DEG motifs typically show the highest pathogenicity, with most scoring above 0.5. Across all SLiM types, the majority of motifs display higher pathogenicity scores than their flanking regions, suggesting that AlphaMissense may be a valuable tool for SLiM detection. b) Cross-validation accuracy distribution of classification of the Decision Tree for Known and PEMs. c) Metrics with the identified PEM motifs. Left panel shows the retention metrics while the right panel shows pathogenic mutations and motifs with pathogenic mutations. d) Benign Mutation Rate for refined PEMs compared to all regex matched motifs.

## Notes

### Competing Interest Statement

The authors have declared no competing interest.

### Summary of Updates

This manuscript has undergone a complete textual revision to improve clarity, readability, and narrative structure. The title has also been updated to better reflect the content. Importantly, all underlying scientific data, analyses, and the core results remain unchanged from the previous version.

